# A zebrafish reporter line reveals immune and neuronal expression of endogenous retrovirus

**DOI:** 10.1101/2021.01.21.427598

**Authors:** Holly A. Rutherford, Amy Clarke, Emily V. Chambers, Jessica J. Petts, Euan G. Carson, Hannah M. Isles, Alejandra Duque-Jaramillo, Stephen A Renshaw, Jean-Pierre Levraud, Noémie Hamilton

## Abstract

Endogenous retroviruses (ERVs) are fossils left in our genome from retrovirus infections of the past. Their sequences are part of every vertebrate genome and their random integrations are thought to have contributed to evolution. Although ERVs are mainly kept silenced by the host genome, they are found activated in multiple disease states such as auto-inflammatory disorders and neurological diseases. What makes defining their role in health and diseases challenging is the numerous copies in mammalian genomes and the lack of tools to study them. In this study, we identified 8 copies of the zebrafish endogenous retrovirus (*zferv*). We created and characterised the first *in vivo* ERV reporter line in any species. Using a combination of live imaging, flow cytometry and single cell RNA sequencing, we mapped *zferv* expression to early T cells and neurons. Thus, this new tool identified tissues expressing ERV in zebrafish, highlighting a potential role of ERV during brain development and strengthening the hypothesis that ERV play a role in immunity and neurological diseases. This transgenic line is therefore a suitable tool to study the function of ERV in health and diseases.

## Introduction

Over 40% of the human genome comprises endogenous transposable elements capable of recombination and disruption of genes and modification of their expression (Lander et al. 2001). Endogenous retroviruses (ERVs) are transposable elements originating from old integrations of retroviruses so successful that they have become part of most vertebrate genomes studied. ERVs replicate autonomously using a copy-and-paste mechanism and although they form a smaller percentage of all retroelements, they still represent 8% of the human genome (Bourque et al. 2018; Lander et al. 2001). The majority of known ERVs have lost their ability to replicate, but those most recently acquired still have intact genomes with the ability to produce viral RNA and particles. However, many of competent ERVs are under strict transcriptional suppression by epigenetic mechanisms (Maksakova, Mager, and Reiss 2008; Rowe et al. 2010; Turelli et al. 2014), protecting the host organisms against potential retroviral insertions and viral activities.

Although immobilised by mutations or transcriptionally repressed, ERVs have a complex relationship with the human genome, which they can regulate by providing cis-regulatory elements to surrounding genes and by lifting their transcriptionally repressed state. Through these mechanisms, it is believed that transposable elements have fuelled some of the necessary genetic changes for evolution (Feschotte 2008; Kunarso et al. 2010). *Syncytin-1*, an ERV envelope gene essential for the vascularisation of the placenta, is at the origin of evolutionary diversification of the placenta (Chuong 2018; Mi et al. 2000). During human embryogenesis, expression of specific ERV families have been associated with cell identity and cell potency in early stem cells (Göke et al. 2015). Additionally, ERV expression has been reported in healthy human tissues such as ovary and testis for ERV-9 (Pi et al. 2004), pancreas (Shiroma et al. 2001), breast (Tavakolian, Goudarzi, and Faghihloo 2019), stomach and small intestine (Okahara et al. 2004). Mainly based on transcriptional studies, the expression of different families of ERV is likely to be extended to more tissues, however their role in tissue development and function remains largely unknown.

ERVs have been linked directly and indirectly to the evolution and functioning of the immune system. Enhancer regions of interferon stimulated genes key to the interferon pathway, such as IRF1 and STAT1, were found introduced and amplified by ERV elements, with the human inflammasome failing to form upon the deletion of a subset of ERVs (Chuong, Elde, and Feschotte 2016). Adaptive immunity also benefits from the presence of ERVs. Indeed, ERV peptide recognition is used in T cell selection to optimise antigen recognition and T cells have a higher sensitivity to exogenous virus infection when presented with ERV peptides during their initial thymic selection (Mandl et al. 2013; Young et al. 2012). The human ERV (HERV) envelope gene contains immunosuppressive domains that reduce the Th1 response during pregnancy and therefore promote foetal development (Knerr et al. 2004; Lokossou et al. 2020). The role of ERVs in our immune system, particularly in the training of our adaptive immunity, can be a double-edge sword as ERVs are linked to a range of different disease states, including autoimmunity.

Aberrant expression of ERVs contributes to multiple pathologies. ERVs are found in abundance in multiple forms of cancers and are considered tumour-promoting factors (extensively reviewed in (Bermejo et al. 2020). The pathology of auto-inflammatory diseases has also been strongly associated with ERVs. Systemic lupus erythematosus (SLE) is an autoimmune disorder with increased level of autoantigen for an ERV (Jorgensen et al. 2014). Recently, one of the murine SLE susceptibility locus was identified as a key suppressor of ERV expression, consolidating the role of ERVs in the pathogenesis of SLE (Treger et al. 2019). A similar disorder is Acardi-Goutieres Syndrome (AGS), a type 1 interferonopathy which resembles congenital cytomegalovirus infection and is caused by mutation in several genes encoding enzymes responsible for nucleic acid metabolism (Crow et al. 2015). Mutations in some of these genes, such as TREX1, MDA5 and ADAR1 trigger aberrant presence of various retroelements and the upregulation of an antiviral immune response (Ahmad et al. 2018; Benitez-Guijarro et al. 2018; Chung et al. 2018; Li et al. 2017; Thomas et al. 2017). Interestingly, anti-reverse transcriptase therapy in AGS patients can decrease the IFN response, highlighting the role of aberrant presence of ERVs in triggering an immune response (G. Rice et al. 2018). Increased expression of ERVs has been found in brains of patient suffering from neurodegenerative diseases such as Motor Neuron Disease (Li et al. 2015) and multiple sclerosis (Johnston et al. 2001; Mameli et al. 2007). Overexpression of a human ERV in neurons was shown to be neurotoxic, suggesting a potential role of ERVs in triggering neuronal toxicity (Li et al. 2015). A direct link to the pathology of these disorders has yet to be made, but nonetheless ERVs appear as strong causal factors for autoimmune and neurological disorders.

Although ERV enrichment has been detected in neurological pathologies, little is known about the function of ERV in healthy tissues. The exact role of ERV in our immune system and brain pathologies has yet to be understood and there is no model system specifically looking at ERV function *in vivo*. Zebrafish is already established as a model to study the immune system, with significant homology with mammals in innate and adaptive immunity (Renshaw and Trede 2012; Trede et al. 2004). The genetic tractability and transparency of the zebrafish embryos have allowed the creation of transgenic reporter lines, some of which have elucidated the role of immune cell behaviour *in vivo* (Renshaw et al. 2006). The tractability of the zebrafish has already been exploited to visualise the expression of the human ERV-9 in oocytes, similarly to human expression (Pi et al. 2004).

In this study, we used the zebrafish as a tractable *in vivo* model to characterise the zebrafish endogenous retrovirus (*zferv*). We identified multiple *zferv* family members, including 2 complete genomes *zferv1a* and *zferv1b*. Using the transparency of the zebrafish larvae we developed a reporter line for *zferv1a* and imaged for the first time ERV activation in healthy tissues *in vivo*. We showed that *zferv1a* is expressed in the thymus and in the brain. Colocalisation analysis and single cell RNA sequencing revealed expression of *zferv1a* specifically in T-cells, suggesting a potential role for ERV in lymphocyte function or development. Brain expression of ZFERV appears to include neuronal expression. This transgenic line can be used as a tool to further investigate the role of ERVs in immunity and in neurological disorders.

## Results

### The zebrafish genome contains multiple endogenous retrovirus integrations

The presence of an ERV in zebrafish, named *zferv*, has been reported by Shen and Steiner while screening a thymic cDNA library (Shen and Steiner 2004) (**Figure 1A**). We searched for related sequences in the most recent reference zebrafish genome (GRCz11, Tü strain) using BLASTN searches, with the original *zferv* sequence as a query. Limiting ourselves to sequences flanked by long terminal repeats (LTRs) on both sides, we retrieved 8 hits scattered in the zebrafish genome, but not the exact *zferv* sequence, possibly because of strain difference (**Figure 1+ Sup. Table 1**). We identified 2 sequences encoding apparently fully functional ERVs, with 91% and 90% identity to *zferv*, that we respectively named *zferv1a* and *zferv1b* (**Figure 1B**). These two ERVs encode almost identical proteins (95 to 97% identity at the amino-acid level) and have LTR promoter region highly homologous (95% identity at the nucleotide level). An additional 6 pseudo *zferv* genes (here called *zferv2* - *zferv7*) were identified (**Figure 1C**), *zferv2* containing a frameshift, *zferv3* and *zferv4* with large deletions and *zferv5, zferv6* and *zferv7* containing a large insertion, all resulting in a degenerated ERV genome.

**Figure 1.**
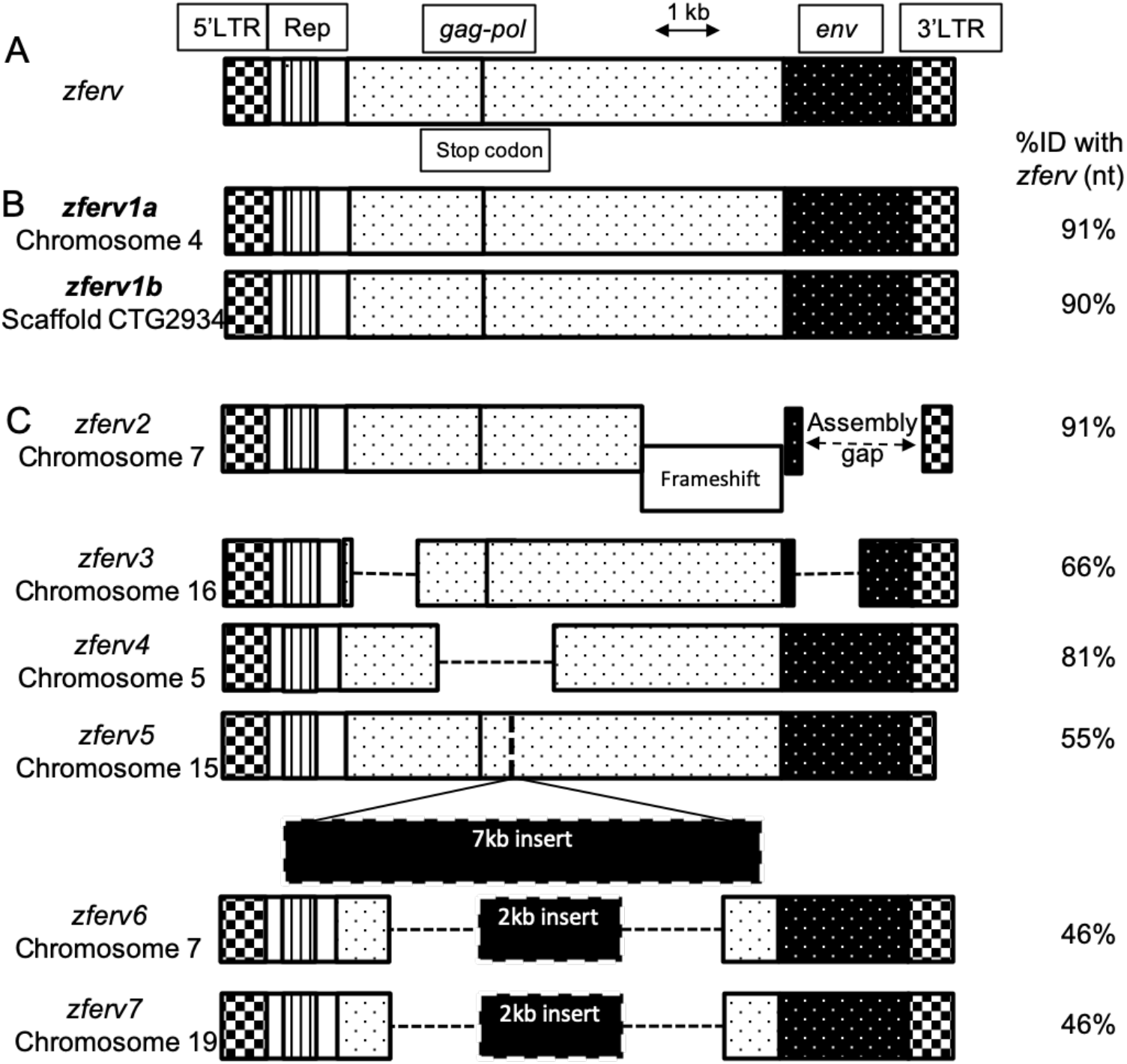
Multiple copies of zferv are present in the zebrafish genome. **A**. Diagram of the original zferv genome found by Steiner et al. used as a reference for nucleotide identity (ID% nt) **B**. Diagram representing the two closest related *zferv* genome found in most recent zebrafish genome GRCz11, named zferv1a and *zferv1b*. **C**. Diagram of 6 pseudo zferv genes with degenerated genome (dotted line represents insertions). 5’-LTR: 5’ –Long terminal repeat, Rep.: Repetitive element, gag-pol: genes encoding the polyprotein and reverse transcriptase, env: envelope gene.

### *zferv1a* is actively expressed in the brain and in the thymus

*zferv* expression was initially found in the thymus at 5 days post fertilisation (dpf) using an RNA probe against the envelope (*env*) gene of the original *zferv* (Shen and Steiner 2004). To verify this observation, the entire *zferv1a* genome was subcloned from a fosmid provided by the Sanger Center and used to create an *in situ* hybridisation (ISH) RNA probe against the envelope gene of *zferv1a*, here called *env1a*. ISH performed on a time course starting from 2 cell-stage embryos until 5days post fertilisation (dpf) confirmed a strong signal in the thymus at 5dpf (**Figure 2A**), similarly to what was previously reported (Shen and Steiner 2004). Individually labelled cells were visible around the thymus following the branchial arches and around the ear (**Figure 2A, black box**). Additionally, we identified a signal at earlier timepoint during gastrulation and a clear signal in the brain, particularly strong at 3dpf (**Figure 2A**). Neuromasts were labelled after 24h exposure to the probe (**Sup. Figure 1**). There was some signal in the thymus using the sense probe signal, suggesting some bidirectional expression of *zferv1a*, as reported for some human ERV (Chiappinelli et al. 2015).

**Figure 2:**
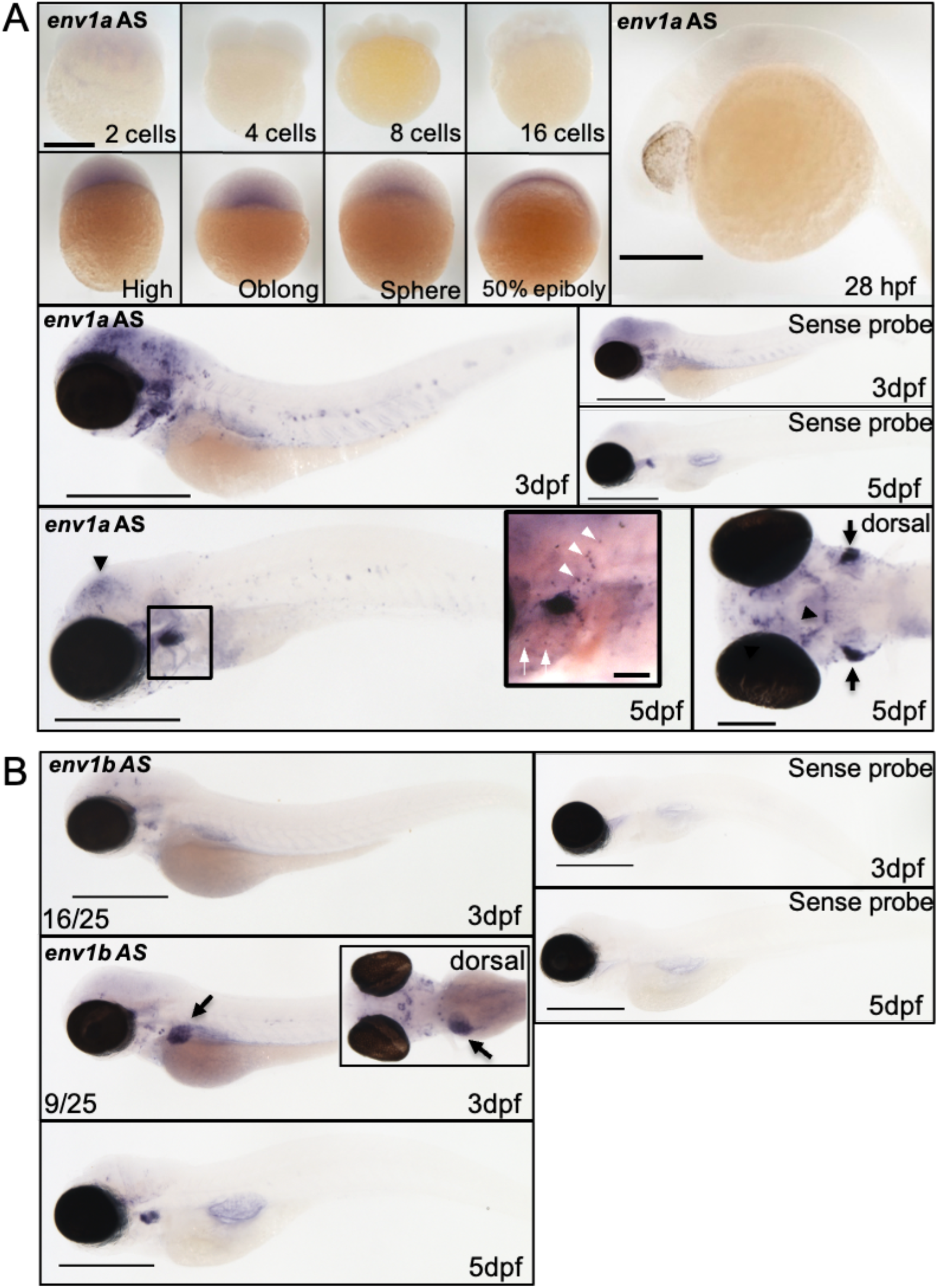
Reporter line for *zferv1a* recapitulates endogenous expression. **A**. Expression of the envelope gene (*env*) of *zferv1a* (called *env1a* here) using an antisense (AS) and sense *in situ* hybridisation RNA probe from 2-cell stage until 5dpf (black arrowheads pointing to brain expression). Scale bar 500μm. Dorsal view of *env1aAS* expression in the brain (black arrowheads) and thymus (Black arrows). Scale bar 200μm. Black box is a zoomed image on the thymus area, highlighting strong expression around the thymus with single positive cells visible in the vicinity of the thymus around the ear (white arrowheads) and alongside the branchial arches (white arrows). Scale bar 70μm. **B**. Expression of the envelope gene (*env*) of *zferv1b* (called *env1b AS* here) by *in situ* hybridization at 3dpf and 5dpf. Liver expression highlighted by black arrows. Scale bar 500μm.

To ensure the specificity of this probe to the *zferv1a* gene, we created a probe against the reverse transcriptase gene of *zferv1a* (*pol1a*) and showed similar labelling of the brain and thymus (**Sup. Figure 2)**. We additionally created a probe against another *zferv* isoform, using specific primers (**Sup. Figure 3**) to the envelope gene of *zferv1b (*called here *env1b)*. Although we still identified a signal in the thymus at 5dpf, we observed a different expression pattern from *env1a* with no brain expression and a strong labelling of the liver in some embryos at 3dpf (**Figure 2B**).

### *zferv1a* reporter line recapitulates endogenous expression

To follow the expression of the zebrafish endogenous retrovirus we identified as *zferv1a*, we took a transgenic approach taking advantage of the promoter activity of retroviral LTR. We used the 5’ viral promoter *ltr5* from *zferv1a* to drive GFP expression by Gateway recombination (**Figure 3A**). Injected embryos showing expression in their thymus were raised and screened in adulthood for germline transmission; one founder was selected to generate the transgenic line bearing the sh533 allele, which was analysed at the F2 generation. Comparable to what was observed by ISH, transgenic embryos (obtained from transgenic mothers) displayed GFP signal at the blastula stage developed a strong GFP signal in the brain by 3dpf (**Figure 3C,D**). GFP signal observed in the eye was caused by the reflective nature of the retina and not by expression of the transgene, as shown using illumination from different laser also highlighting the eye and in situ hybridisation labelling with the env1a probe in a 5dpf embryo with pigment partially removed in the eye using a tyrosinase crispant (**Sup. Figure 4**). A weak signal in the thymus can be observed at 3dpf (**Figure 3C** white arrowhead) as the thymus starts to develop. At 5dpf, a strong signal was observed in the thymus (**Figure 3E,F**), comparable to that observed by ISH (**Figure 1A,B**). Consistent with ISH, our reporter line also labelled the optic tectum region of the brain and the spinal cord at 5dpf, with axons labelled in individual neurons (**Figure 3G,H**). GFP signal was also found in neuromasts (**Figure 3E, white arrows**), which we also observed in a small proportion of 5dpf embryos by ISH with longer development time (**Sup. Figure 1**). Transmission rate of 50% to F3 embryos of the brain signal was variable depending which F2 animal was used, including complete and partial brain labelling, whereas the thymic expression remaining constant throughout the generations.

**Figure 3:**
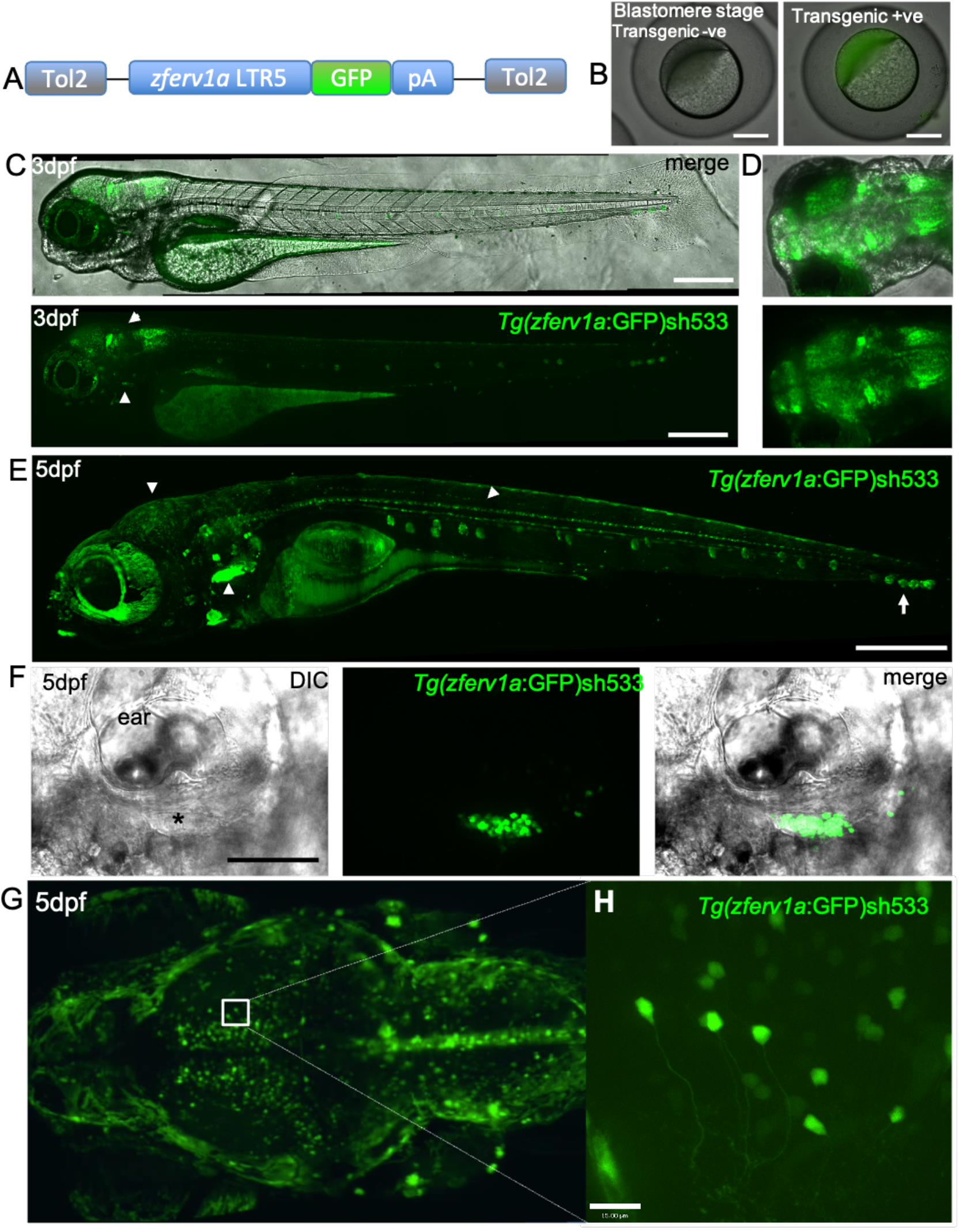
Reporter line for *zferv1a* recapitulates endogenous expression. **A**. Diagram of gateway construct used to create the *Tg(zferv1a:GFP)* reporter line using the pDestCryCFP tol2 backbone. **B**. Representative image at blastula stage in non-transgenic and *Tg(zferv1a:GFP:pA)sh533* transgenic embryos. **C**. Lateral view of DIC and GFP high resolution images of a 3dpf *Tg(zferv1a:GFP:pA)sh533* embryos, displaying high expression of the transgene in the brain (white arrow) and the start of a signal in the thymus (white arrowheads). Scale bar 400μm. **D**. Dorsal view of the brain at 3dpf, merged with with brightfield image and GFP only. **E**. High resolution image of a 5dpf *Tg(zferv1a:GFP:pA)sh533* embryos, displaying high expression of the transgene in the thymus (white arrowheads), brain and spinal cord (white arrowheads). Note signal in neuromast alongside the trunk (white arrow). Scale bar 500μm. **F**. Single slice DIC and confocal GFP fluorescence merged image of the thymus (black star) situated underneath the ear in 5dpf embryos and the signal from the *Tg(zferv1a:GFP)sh533* reporter. Scale bar 100μm. **G**. Dorsal view of a maximum projection of the brain from 5 dpf *Tg(zferv1a:GFP*) embryo acquired with a the lightsheet microscope. **H**. High resolution single slice image from the optic tectum of *Tg(zferv1a:GFP)*. Scale bar 15 μm.

This transgenic line was not the first attempt to create a fluorescent reporter for *zferv1a*. Our initial reporter line was made using both 5’ (*ltr5*) and 3’ (*ltr3*) promoter regions in order to resemble the genome structure of an ERV. This line, called Tg(*ltr5:*GFP*:ltr3*)*sh500*, contained the CryCFP eye marker to help with selection of positive embryos. Although outcrossed F1 showed normal transmission rate (50% of positive embryos in F2, with CFP in lens and GFP in thymus), F2 transmission rate became unstable. Selected F2 positive embryos grown to adulthood failed to transmit the same signal pattern, with often reduced to complete loss of thymus labelling and random brain expression pattern despite transmission of the CFP lens phenotype at expected mendelian ratios. We also observed a dampening of the GFP signal in F3 and complete loss in F4. This suggested that the transgene was behaving as a transposable element and was ultimately silenced by the host or lost in the germline. This line was abandoned, and we continued our study with the *sh533* line.

### Zebrafish endogenous retrovirus *zferv1a* expression is restricted to lymphoid cells

To further study the nature of the cells labelled in the thymus in the *sh533* line, we took advantage of the transparency and ease of access of the thymic tissue to perform live imaging. Cell tracking of timelapse imaging of the *sh533* line revealed a dynamic behaviour of GFP labelled cells, approaching and exiting the thymus following the ear and the branchial arches (**Figure 4A, Movie 1**), as previously described for thymocytes (Dee et al. 2016; Kissa et al. 2008). This thymic signal persisted beyond embryonic stages and was still visible in juveniles at 25dpf (**Figure 4B**). To analyse the origin of the *zferv1a* cell population, we dissociated 3 head kidneys from Tg(*zferv1a*:GFP)*sh533* and non-transgenic control adults and used size and granularity scatter by flow cytometry to distinguish between the various haematopoietic lineages (**Figure 4C**). According to these criteria and using a back-gating method for quantification (**Figure 5D,E, Sup. Figure 5**), we found that GFP positive cells have a typical lymphocyte profile (**Figure 4F**).

**Figure 4:**
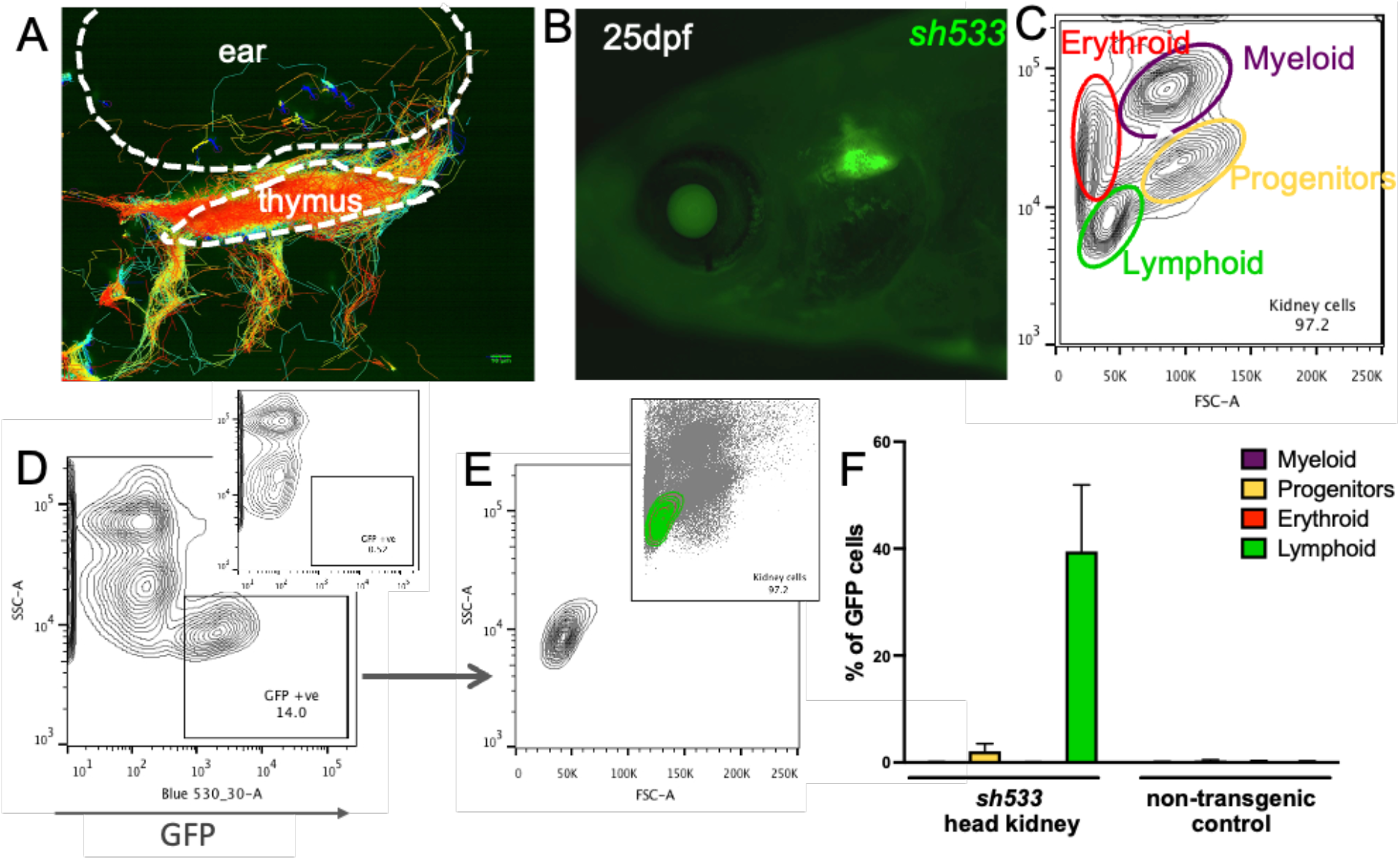
*zferv1a* is expressed in cells from the lymphoid lineage in adult haematopoietic tissues. **A**. Cell tracking reveals dynamic behaviour of entry and exit of thymus by GFP positive cells in Tg(zferv1a:GFP)*sh533*. Whole stack analysis of cell movement by using TrackMate on FiJi. Scale bar 10μm. **B**. Expression of *Tg(zferv1a:GFP)sh533* in a 25dpf zebrafish highlighting the triangular shaped thymus (white arrowhead). **C**. FSC/SSC plot of a whole kidney marrow separating different haematopoietic lineages by size and granularity. **D**. Side scatter (SSC) and GFP plot from single cell suspension of a whole kidney marrow from an adult *Tg(zferv1a:GFP)* with gate selecting GFP positive cells. Inset plot showing the distribution of cells from a non-transgenic adult whole kidney marrow. **E**. Forward and side scatter (FSC/SSC) plot of selected GFP positive cells from panel D. Inset plot showing the location of the GFP positive cells in the haematopoietic lineage FSC/SSC plot. **F**. Quantification of GFP positive cells in myeloid, erythroid, lymphoid lineages and progenitors, as defined in the dot plot D, from dissected sh533 and non-transgenic control adult head kidney (n=3).

**Figure 5:**
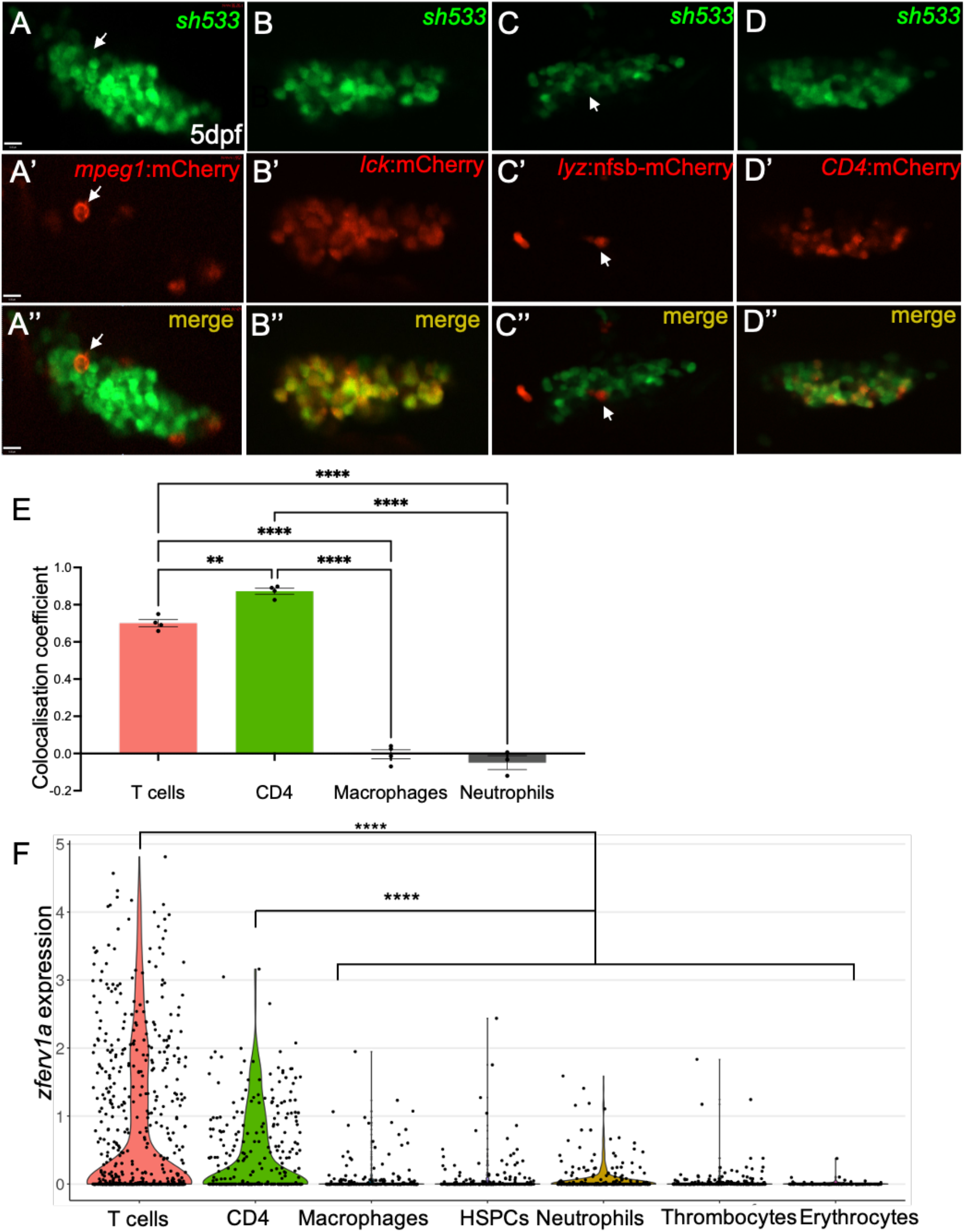
Immune expression of *zferv1a* is restricted to CD4 and T-cells. **A-D**. Single slice confocal image of the thymus from double transgenic *Tg(zferv1a:*GFP*)sh533 x Tg(mpeg1:*mCherryCAAX)*sh378* labelling macrophage membrane (A-A’’-white arrow pointing to a single macrophage), x *Tg(lck:*mCherry) labelling T cells (B-B’’), *x Tg(lyz:nfsb-*mCherry)*sh260* labelling neutrophils (C-C’’-white arrow pointing to a single neutrophil), and x *Tg(CD4:*mCherryCAAX) labelling CD4 cells (D-D’’) with merge image used for colocalisation analysis. **E**. Histogram plot of ImageJ-generated colocalisation coefficient of *Tg(zferv1a:*GFP*)sh533* crossed to *Tg(mpeg1:*mCherryCAAX)*sh378* for macrophages, *Tg(lyz:nfsb-*mCherry)*sh260* for neutrophils, *Tg(CD4:*mCherry) for CD4+ cells and *Tg(lck:*mCherry) for T cells (one-way ANOVA, n=4). **F**. Violin plot of single-cell RNA sequencing datasets MTAB-5530 and E-MTAB-4617 from zebrafish adult whole kidney marrow and spleen showing expression of *zferv1a* (*ENSDARG00000110878*) in multiple haematopoietic lineages. Mann-Whitney with paired Wilcox Test. p = 1.95 × 10^−11^ for CD4 versus the other clusters and p = 1.97 × 10^−45^ for T cells versus the other clusters.

To further corroborate the lymphoid identity of *zferv1a* expressing cells, we performed *in vivo* colocalisation analysis in 5dpf embryos by crossing the Tg(*zferv1*a:GFP)*sh533* line to different reporter lines labelling a wide spectrum of immune cell populations. We crossed F1 Tg(*zferv1a*:GFP)*sh533* to reporter lines labelling lymphoid cells, such as Tg(*lck*:mCherry), labelling T cells (Langenau et al. 2004) but also NK and innate lymphoid cells (Hernández et al. 2018), or Tg(*CD4*:mCherry), labelling CD4+ T cells and some macrophage/dendritic cells (Dee et al. 2016) and showed significant colocalisation between GFP and mCherry cells by colocalisation analysis (**Figure 5B,D,E)**. By contrast, crossing of the Tg(*zferv1a*:GFP)*sh533* to macrophage Tg(*mpeg1*:mCherryCAAX)*sh378* (**Figure 5A**) or neutrophil *Tg(lyz:nfsb-*mCherry)*sh260* (**Figure 5C**) (Buchan et al. 2019) reporter lines did not reveal significant colocalisation **(Figure 5E)**. To validate these results, we used single cell RNA sequencing datasets available on zebrafish blood cells (MTAB-5530 and E-MTAB-4617) as a robust assay to analyse gene expression in specific cell type and tissues. Using the extensive gene expression dataset available, we mapped the expression of the entire genome of *zferv1a* (*ENSDARG00000110878)* in erythrocytes, haematopoietic stem and progenitor cells (HSPCs), macrophages, neutrophils, CD4, T cells and thrombocytes. Only sporadic expression was observed in all cell lineages except for T cell subsets, CD4+and T cells, where significant expression of *env* was detected (**Figure 5F**). Altogether, these data are consistent with immune expression of *zferv1a* being restricted to T cells.

### Modulation of *zferv1a* expression

To investigate whether *zferv1a* can affect the development of the tissues where it is expressed, we performed an overexpression assay. We injected the vector containing *zferv1a* alongside the same vector containing the eGFP cassette to use as a positive control. Each plasmid was injected in equimolar amount or GFP alone and the expression of zferv1a and GFP was measured by qPCR. Overexpression of GFP resulted in a high level of GFP transcript detected by qPCR, however we failed to observe an increase in zferv1a expression when injected (**Sup. Figure 6**). We then checked for induction of *zferv1a* (*ENSDARG00000110878*) in our datasets of larvae stimulated with type I interferon or with chikungunya (CHIKV) Virus ((Levraud et al. 2019); bioproject accession# PRJNA531581). No expression was detected in the dataset of type I IFN injected larvae (3dpf). Expression was detected in the dataset of CHIKV-infected fish (4dpf). However, while CHIKV infection triggers a very strong interferon response, with 1000-fold induction of some ISGs, there was only a statistically non-significant 1.7-fold increase for *zferv1a*.

## DISCUSSION

We describe here a novel reporter line in zebrafish to follow expression of an endogenous retrovirus in real time *in vivo*. Our results validate and refine previously published expression data of *zferv* in the thymus and specifically in the lymphoid lineage. We identified a close relative to *zferv*, here named *zferv1a*, and showed transcriptional activity mostly restricted to T lymphocytes as previously reported (Frazer et al. 2012; Shen and Steiner 2004), but also observed in neurons.

### Stability of engineered-ERV transgene

Using a transgenic approach, we imaged *in vivo* in real time the behaviour of an ERV for the first time. Transgenesis entails random insertion of the transgene in the genome, which in some cases can disrupt neighbouring gene expression or induce silencing of the transgene (Akitake et al. 2011). Our initial ERV transgene Tg(*ltr5*:GFP:*ltr3*)*sh500* was engineered to resemble the ERV structure and keep both LTR sequences. However, this line started to show signs of genetic instability and silencing from the F2 generation. It is possible that the location of the insertion triggered silencing of the transgene, however, we suspect that due to the similarity to a retrovirus genome, the transgene was targeted by epigenetic mechanisms specific to ERV. Indeed, transcriptional repression of ERV has been extensively studied and key histone and DNA methylation modifications identified (Maksakova et al. 2008; Rowe et al. 2010; Turelli et al. 2014), which could be targeting the LTR promoter we used. Previous work on *zferv* identified amplification of the *zferv* locus initially in T cells from an acute lymphoblastic leukemia zebrafish model and, although at a lesser rate, also in normal T cells (Frazer et al. 2012). We suspect that genomic amplification of the initial transgene *ltr5*:GFP:*ltr3* would be facilitated in the initial sh500 transgenic line as the transgene would be subjected to the transposase activity of other functional ERVs. This in turn might have activated targeting and silencing of the transgene within a generation. Either or both of those mechanisms might explain why we still observed some mild (neuron-specific) silencing phenotypes in the second transgenic line using a less recognisable ERV structure, with the 3’LTR sequence replaced by a normal polyA tail in Tg(*ltr5*:GFP:*pA*)*sh533*.

### Role of ERVs in T lymphocytes

Despite stability issues of the transgene, we consistently observed the signature expression of *zferv* in lymphocytes, already identified in zebrafish transcriptional studies by others (Frazer et al. 2012; Shen and Steiner 2004). The role of ERV expression in T cells is still under investigation. Each step of T cell differentiation is accompanied by a sequence of wide transcriptional changes that allows 1) commitment to the T cell lineage, 2) T cell receptor (TCR) rearrangement and 3) positive or negative selection (Mingueneau et al. 2013). It is possible therefore that transcriptional de-repression of neighbouring genes allows ERV to be expressed during those stages, which would explain such a strong expression during those early phases of T cell development.

Whether ERV expression is merely a side effect of neighbouring gene activation or plays a crucial role during T cell development is an important question to answer. ERV do confer advantages to our adaptive immune system, with important implications for infectious and cancer therapeutic approaches. T cells that have been presented with ERV peptides during their initial thymic selection have a higher sensitivity to exogenous retrovirus infection, conferring an advantage in fighting HIV (Young et al. 2012). And very recently, T cell populations were found with the ability to recognise the antigen of multiple HERV in patients with haematological cancers, with HERV-T cell less present in healthy donors (Saini et al. 2020). This finding might lead to selective therapies using HERV as a target to kill tumour cells.

### Relevance of neuronal expression of ERV for human studies

We are providing evidence for a member of the ERV family to be expressed in the healthy developing brain of a vertebrate. Reporter expression suggests that this neuronal expression might be more easily silenced than in T cells; the relevance of this finding for endogenous *zferv* expression, and its physiological significance, remain to be investigated. The role of endogenous retrovirus expression in developing neurons deserves further investigation as the potential role of ERVs in neurodegenerative and neurological diseases is a subject under intense study. The implications of aberrant presence of ERVs in the brain in human neurological diseases are actively being investigated, with the human HERV-K found upregulated in cortical neurons of ALS and MS patients (ref), with currently active clinical trials targeting ERV with anti-retroviral therapies for ALS, MS and the autoinflammatory disease AGS (Gold et al. 2018, 2019; G. I. Rice et al. 2018).

Our finding in embryos and adults suggests a role of ERV in young neurons, potentially highlighting a role for ERV in neuron development and maturation. There is evidence that other types of retroelements are expressed in neurons where they might function as regulators of progenitor differentiation. Indeed, brain expression of other retroelements has previously been reported, mainly focused on the family of non-LTR element, the Long Interspersed Nuclear Element L1, comprising the largest percentage of the human genome of all RE accounting for 17% of all genomic DNA (Scott et al. 1987). In the brain, L1 elements are de-repressed to allow neuronal precursor cells differentiation and contribute to individual somatic mosaicism in human (Coufal et al. 2009) and mouse (Muotri et al. 2005). The latest study also used an engineered reporter approach by tagging GFP to the retroelement L1 (Muotri et al. 2005). Although L1 elements are not related to ERV which is the subject of our study, it was recently discovered that additional groups of RE are enriched in specific regions of the human brain, LTR elements in the cerebellum in particular (Bogu et al. 2019). With the advances in single cell sequencing, it is possible that expression of different groups of ERV, specifically human ERVs, will also be identified in the brain to begin a complete understanding of the presence and role of ERV in brain development and diseases (Evans and Ann 2020).

### Conclusions and future work

The zebrafish has been successfully used to model human neurodegenerative and neurodevelopmental disorders. Early timepoint of brain development, particularly difficult to study in mammals, are studied with ease in the zebrafish due to its transparency. As such, zebrafish models of neurodevelopmental autoinflammatory disorders (Hamilton et al. 2020; Haud et al. 2011; Kasher et al. 2015) or neurodegenerative disease such as Multiple Sclerosis (Kulkarni et al. 2017) or ALS (Da Costa et al. 2014) can be combined to our *zferv1a* reporter line to elucidate the role of ERV in the development of the disease.

The first zebrafish reporter line for an ERV described in our study provides a tool to understand how certain pathological states activates ERV expression in specific cell lineages, using a real time *in vivo* approach.

## METHODS

### Zebrafish husbandry

We used the following transgenic lines: Tg(*mpeg1*:mCherryCAAX)*sh378* labelling the membrane of macrophages and microglia (Bojarczuk et al. 2016), Tg(*lck*:mCherry), labelling T cells (Langenau et al. 2004) but also NK (Hernández et al. 2018), Tg(*CD4*:mCherry) labelling CD4+ T cells and some macrophage/dendritic cells (Dee et al. 2016), the *Tg(lyz:nfsb-*mCherry)*sh260* for neutrophils (Buchan et al. 2019).

All zebrafish were raised in the Bateson Centre at the University of Sheffield in UK Home Office approved aquaria and maintained following standard protocols(Nüsslein-Volhard and Dham 2002). Tanks were maintained at 28°C with a continuous re-circulating water supply and a daily light/dark cycle of 14/10 hours. All procedures were performed on embryos less than 5.2 dpf which were therefore outside of the Animals (Scientific Procedures) Act, to standards set under AWERB (Animal Welfare and Ethical Review

Bodies) and UK Home Office–approved protocols.

### Identification of *zferv* genomes

We used the published full *zferv* nucleotide sequence (AF503912) as a query for a nucleotide BLAST search on zebrafish genome assembly on the Ensembl website (http://www.ensembl.org/Danio_rerio/Tools/Blast?db=core), using the most recent reference genome (GRCz11, as of July 2017), choosing the “BLAT” option favouring highly similar sequences, and disabling the filter for species-specific repeats. We selected only hits with homology found in both the *gag*/*pol* and the *env* regions. Subsequent sequence alignments of LTRs, repetitive element, and open reading frames, and annotations were performed using DNA strider 2.0.

### *zferv1a* full genome cloning

The full zferv1a sequence was subcloned from fosmid 1930h03 (Sanger Center, Cambridge, UK) predicted to contain a ∼37kb of chromosome 4 encompassing the sequence, according the annotations of the zv9 version of the zebrafish genome (Ensembl.org). The sequence, plus ∼100 bases of flanking genomic sequence on each side, was amplified using primers 5’-ccctgctcattcaacaccatac-3’ and 5’-cccgtctgtgaattaccaagc-3’, and cloned into PCR4-TOPO (Thermofisher).

### RNA in situ hybridisation

The *env1a* probe was made from the plasmid containing the full *zferv1a* genome with the following primers: TGGGGATCCATGAATAAAATAAACAAATTGG, BamH1_env1a_fwd and CCACCCGGGCACCATATCCAATAGTTCCTCC, SmaI_env1a_rev generating a 1935bp (env1a) fragment. The *pol1a* probe was made from the same plasmid using TGGGGATCCGGCAGCAGACGCCGCTGCTA, BamH1_pol1a_fwd, TGGTCTAGACCTCAGGCTCCTCAGTGTCT, Xba1_pol1a_rev generating a 1295bp fragment. The *env1b* probe was made from cDNA using TGGGGATCCatgaATATAAACAAATTGGTGG, BamH1_env1b_fwd, CCACCCGGGCAGAACGCTATAGTCAGTGCTC, SmaI_env1b_rev generating a 635bp (env1a) fragment. Fragments were subsequently cloned into Zero Blunt™ TOPO™ vector (Invitrogen) and reverse transcribed into RNA using the DIG labelling kit (Roche) (SP6 enzyme for antisense probe and T7 for sense probe). Whole mount in-situ hybridisation on wild type (WT) nacre zebrafish was performed to identify the spatial and temporal pattern of *zferv1a* expression using the envelope (*env*) probe env1a and the reverse transcriptase probe *pol1a* and from *zferv1b* using the envelope probe *env1b*. We fixed WT embryos from the *nacre* strain at different stages and followed in situ hybridisation protocol as previously described (Thisse and Thisse 2008).

### *zferv1a* and *lck:mCherry* transgenic lines generation

LTR5 and LTR3 regions of *zferv1a* were cloned into p5E and p3E Gateway entry vectors respectively using the following primers: TGTGGATCCTCTCTTTCGAGATCAAGAGAGGG BamH1_LTR3’_fwd, ACACTCGAGGCTGCACCTTGTTGGAAAATGG Xho1_LTR3’_rev, TGTGGTACCCCTCTCTCTTGTAAAGGTTGAGGG kpn1_LTR5’_fwd, ACACCGCGGTTTAATAATGGTTTTGTCTCCC sacII_LTR5’_rev. LR reaction were performed into the pDestCRY-CFP vector and injected into 1 cell stage zebrafish embryos to create the following transgenic lines: *Tg(LTR5:GFP:LTR3)sh500*, and *Tg(LTR5:GFP:pA)sh533*, subsequently called *Tg(zferv1a:*GFP*)sh533*.

The already described *lck* promoter (Langenau et al. 2004) was used to create the transgene *lck*:mCherry in the pDestCRYmCherry vector by Gateway recombination. Stable lines were created by injecting these constructs with Tol2 mRNA in one-cell stage embryos and selecting founders using the red eye marker.

### Tyrosinase CRISPR/Cas9 crispant generation

Synthetic SygRNA (crRNA and tracrRNA) (Merck) in combination with cas9 nuclease protein (Merck) was used for gene editing. Transactivating RNAs (tracrRNA) and gene-specific CRISPR RNAs (crRNA) were resuspended to a concentration of 20 μM in nuclease-free water containing 10 mM Tris–HCl pH 8. SygRNA complexes were assembled on ice immediately before use using a 1:1:1 ratio of crRNA:tracrRNA:Cas9 protein. Gene-specific crRNAs were designed using the online tool CHOPCHOP (http://chopchop.cbu.uib.no/). Tyrosinase crRNA tyr: GGACTGGAGGACTTCTGGGG(AGG)(Isles et al. 2021).

### Flow cytometry analysis

The head kidney of 3 adult *Tg(zferv1a:*GFP*)sh533* and 3 wild type *nacre* fish were dissected and collected into cold Leibovitz 15 media supplemented with 20% FBS and 0.5mM EDTA. The tissue was pipetted up and down to create a single cell suspension and kept on ice. Negative gate was set up using cells from *nacre* animals with the Blue530 detector on an FACSARIA IIu flow cytometry machine and all analysis done using the FlowJo software. Gating was performed as described in supplementary figure 5 using the GFP positive population back gated onto the different cell type population: monocytes, progenitors, T cells and erythrocytes.

### Colocalisation imaging and analysis

*Tg(zferv1a:*GFP*)sh533* was crossed to Tg(*lck*:mCherry), Tg(*CD4*:mCherry), *Tg(lyz:nfsb-*mCherry)*sh260*, Tg(*mpeg1*:mCherry:CAAX)*sh378* and double positive animals were selected and imaged at 5dpf using the Perkin Elmer Spinning Disk system for 3 hours. Whole Z-stacks were imported into FiJi to perform colocalisation analysis using the Colocalisation Threshold Pluggin. Whole Z-ztack at 3 different time points during the timelapse in 4 different larvae were used with the same threshold values to determine the colocalisation coefficient by Pearson’s correlation.

### Single Cell RNA sequencing dataset analysis

Expression of *zferv1a (ENSDARG00000110878*) in different haematopoietic lineages was obtained from single cell RNA sequencing databases of 7 different transgenic lines each labelling a different lineage (Athanasiadis et al. 2017). Single-cell RNA-Sequencing raw counts files of zebrafish blood cells were obtained from the Single Cell Expression Atlas (https://www.ebi.ac.uk/gxa/sc/home, accession codes: E-MTAB-5530 and E-MTAB-4617). The data was then processed and analysed using the Seurat software package for R (Hao et al. 2021; Satija et al. 2015). The data was selected to have a minimum of 200 genes in each cell and then merged and normalised. The Seurat wrapper fastMNN was used to correct for batch effects and UMAP analysis was performed on the resulting combined dataset which resulted in 9 clusters. The function FindAllMarkers was used to find the differentially expressed genes in each cluster and these were compared to marker genes published in the studies to assign cluster identity. Expression distributions across clusters were obtained for ENSDARG00000110878. Differential expression of CD4 and T cells versus all cells (Average log2 fold change = 1.98, adjusted p.value - 5.31e-81). Statistical differential expression was performed in Seurat between each cluster with Mann-Whitney test followed by a Kruskal-Wallis multiple testing correction.

### *zferv1a* overexpression

The plasmid containing the full *zferv1a* gene was injected in equimolar concentration alongside a plasmid containing GFP into 1-cell stage zebrafish embryos, using GFP only injected embryos as control. RNA was extracted from 30hpf GFP positive embryos from both groups using Trizol and cDNA was synthesised as previously described (Hamilton et al. 2020). qPCR was performed to measure level of expression of GFP and *zferv1a* normalised to the *ef1a* reference gene using the following primers: egfpforward: CCATCTTCTTCAAGGACGAC, egfpreverse: CGTTGTGGCTGTTGTAGTTG, ZFERVpol-1S: GCTAGGACATCCCATTGTGT, ZFERVpol-2A: GGGAATGTGTTCTGGTGTCT, Ef1a-1S :GCTGATCGTTGGAGTCAACA, Ef1a-2A: ACAGACTTGACCTCAGTGGT.

### Statistical analysis

All statistical analysis were performed in GraphPad Prism where data was entered using a grouped table for ANOVA analysis (more than 2 samples or 2 variables). Sample distribution was assessed using frequency of distribution analysis. n (experimental unit) number stated for each experiment in figure legends. p values are indicated and a star system is used instead for graph with multiple comparisons: *=p<0.05, **=p<0.01, ***=p<0.001, ****=p<0.0001. Following the recommendation of the American Statistical Association we do not associate a specific p value with significance (Wasserstein, Schirm, and Lazar 2019).

## Funding

This work has been supported by a European Leukodystrophy Association fellowship (ELA 2016-012F4) to NH, an MRC Programme Grant (MR/M004864/1) to SAR and JPL is supported by Agence Nationale de la Recherche (grant ANR-16-CE20-0002-03). Imaging was carried out in the Wolfson Light Microscopy Facility, supported by an MRC grant (G0700091) and a Wellcome Trust grant (GR077544AIA).

## Conflict of Interest Statement

The authors declare that the research was conducted in the absence of any commercial or financial relationships that could be construed as a potential conflict of interest.

## Acknowledgements

We thank the flow cytometry facility at the University of Sheffield, The Light Microscopy Facility and the Zebrafish facility staff at The Bateson Centre for their technical help. We thank Adam Hurlstone (The University of Manchester) for supplying the Tg(*lck*:mCherry) transgenic line, Ana Cvejic for the immune single cell sequencing datset and Angela Ciuffi from Institute of Microbiology (IMUL), Lausanne University Hospital and University of Lausanne for supervising ADJ.

## Author contributions

NH and SAR acquired funding and designed the study; NH, AC, HAR, JJP, HMI, EGC, JPL, AC collected and analysed data. EVC performed all bioinformatic work. ADJ created crucial reagent. NH wrote the manuscript; NH, SAR, JPL revised the manuscript.

---

**Supplementary Table 1:** Coordinates of the *zferv1-7* genomic segments

| ERV name | chromosome (scaffold) | orientation | 5' limit | 3' limit |
| --- | --- | --- | --- | --- |
| *zfer*v1a | 4 | reverse | 62,546,461 | 62,557,677 |
| *zfer*v1b | KN149691.1 - NW_001884427.4 | forward | 211.988 | 223.087 |
| *zfer*v2 | 7 | reverse | 19,379,508 | 19,389,176 |
| *zfer*v3 | 16 | reverse | 9,986,975 | 9,996,169 |
| *zfer*v4 | 5 - NW_018394575 | reverse | 168.525 | 178.11 |
| *zfer*v5 | 15 | forward | 44,144,056 | 44,162,279 |
| *zfer*v6 | 7 - NW_018394672 | reverse | 116.062 | 124.356 |
| *zfer*v7 | 19 | forward | 7,960,461 | 7,968,734 |

**Supplementary Figure 1:**
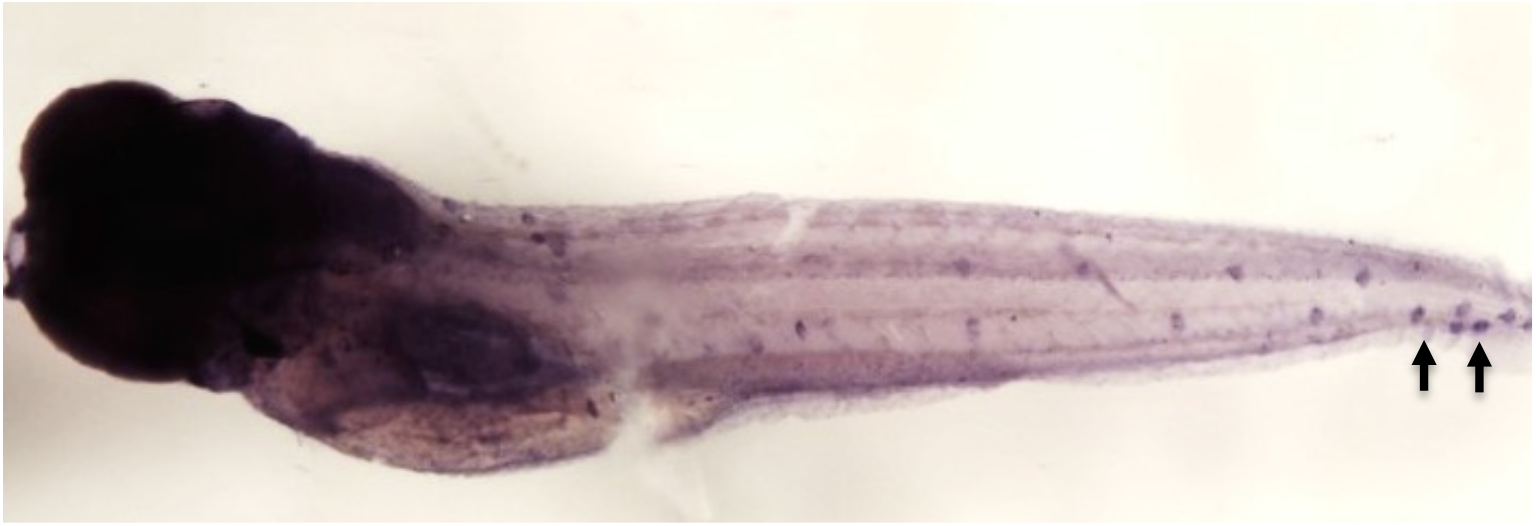
**Long development of the** *env* probe from *zferv1a* by ISH revealed labelling of neuromasts (Black arrows).

**Supplemental Figure 2:**
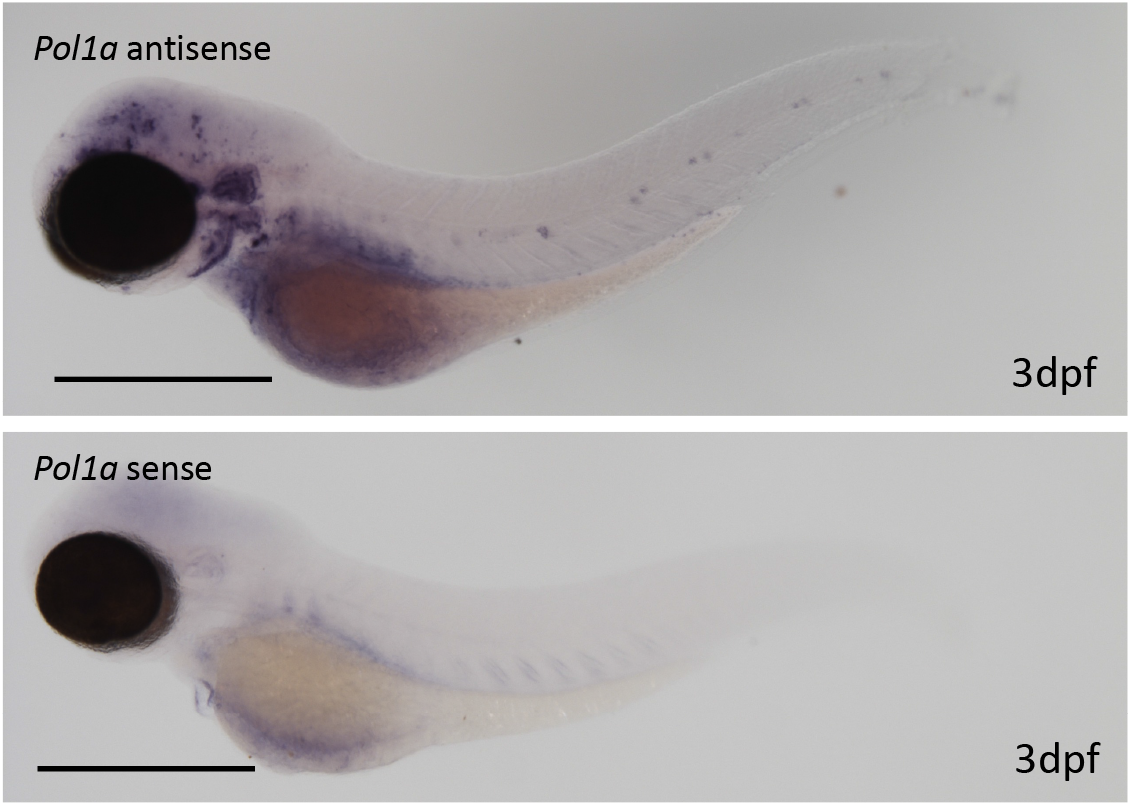
Expression of the reverse transcriptase gene (*pol*) of *zferv1a* (called *pol1a* here) by *in situ* hybridisation at 3dpf. Scale bar 500μm.

**Supplementary Figure 3:**
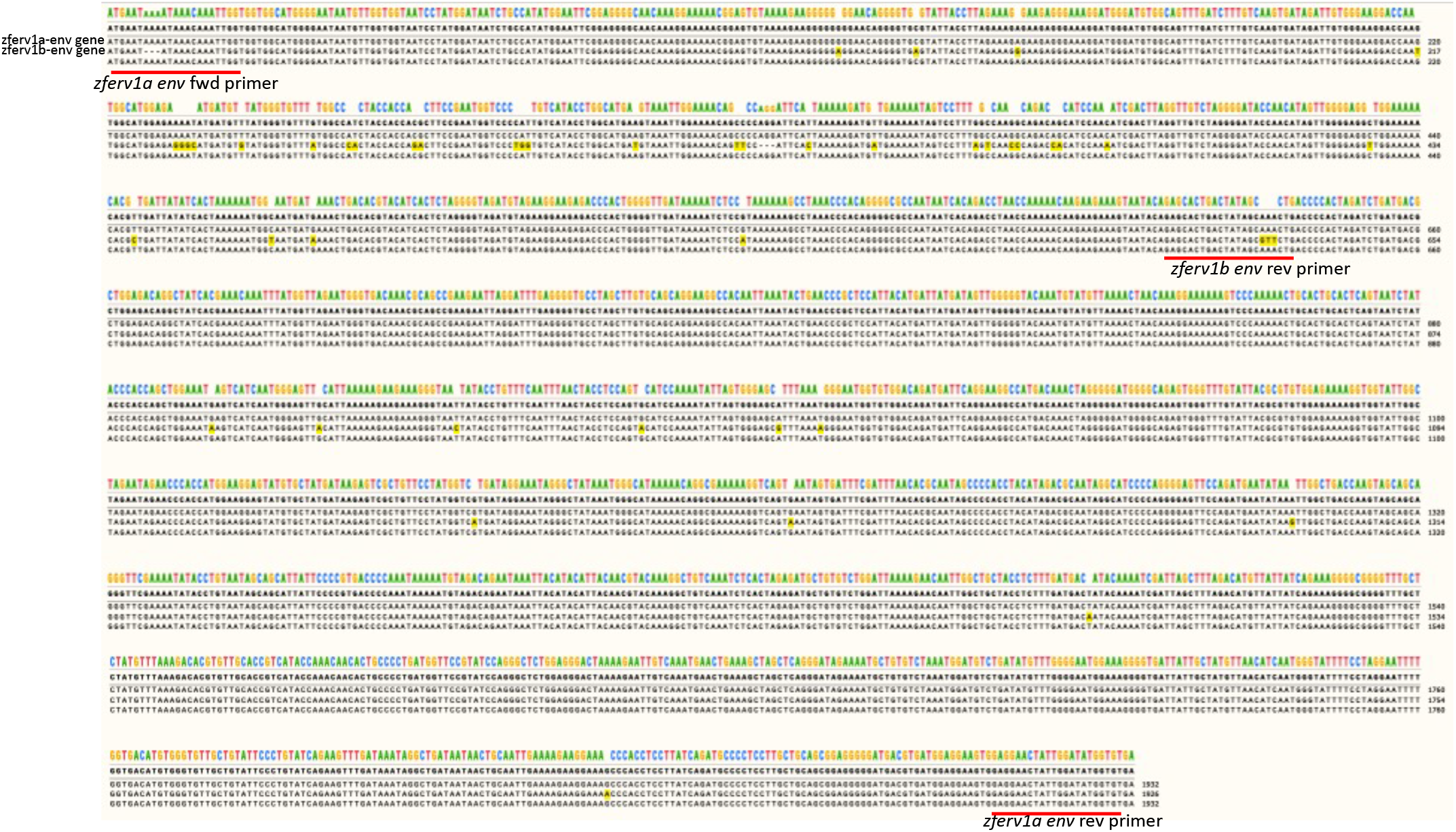
Alignment of the envelope gene from zferv1a and zferv1b to highlight to regions with significant differences between the two genes. Primers used to design the probes are underlined in red.

**Supplementary Figure 4:**
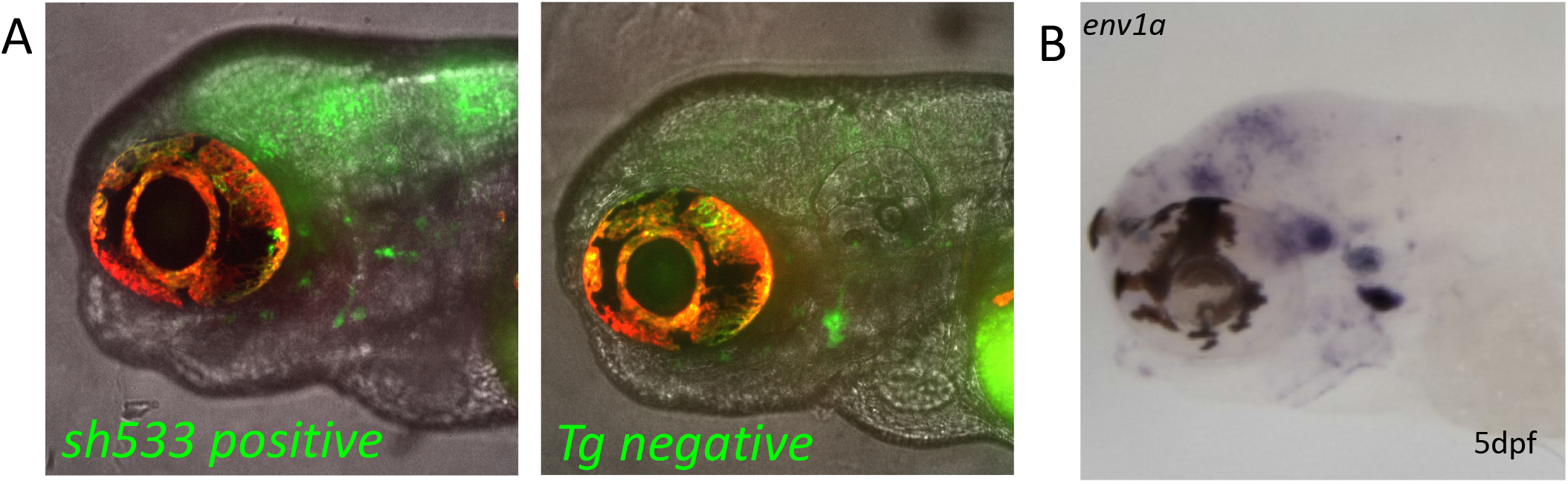
**A**. *sh533* positive and negative embryo illuminated with the GFP and mCherry laser to highlight the similar light reflection in the lenses that act as mirrors. **B. ISH with RNA probe env1a in a 5dpf embryos injected with a crRNA against the tyrosinase gene generating a mosaic pattern of**

**Supplementary Figure 5:**
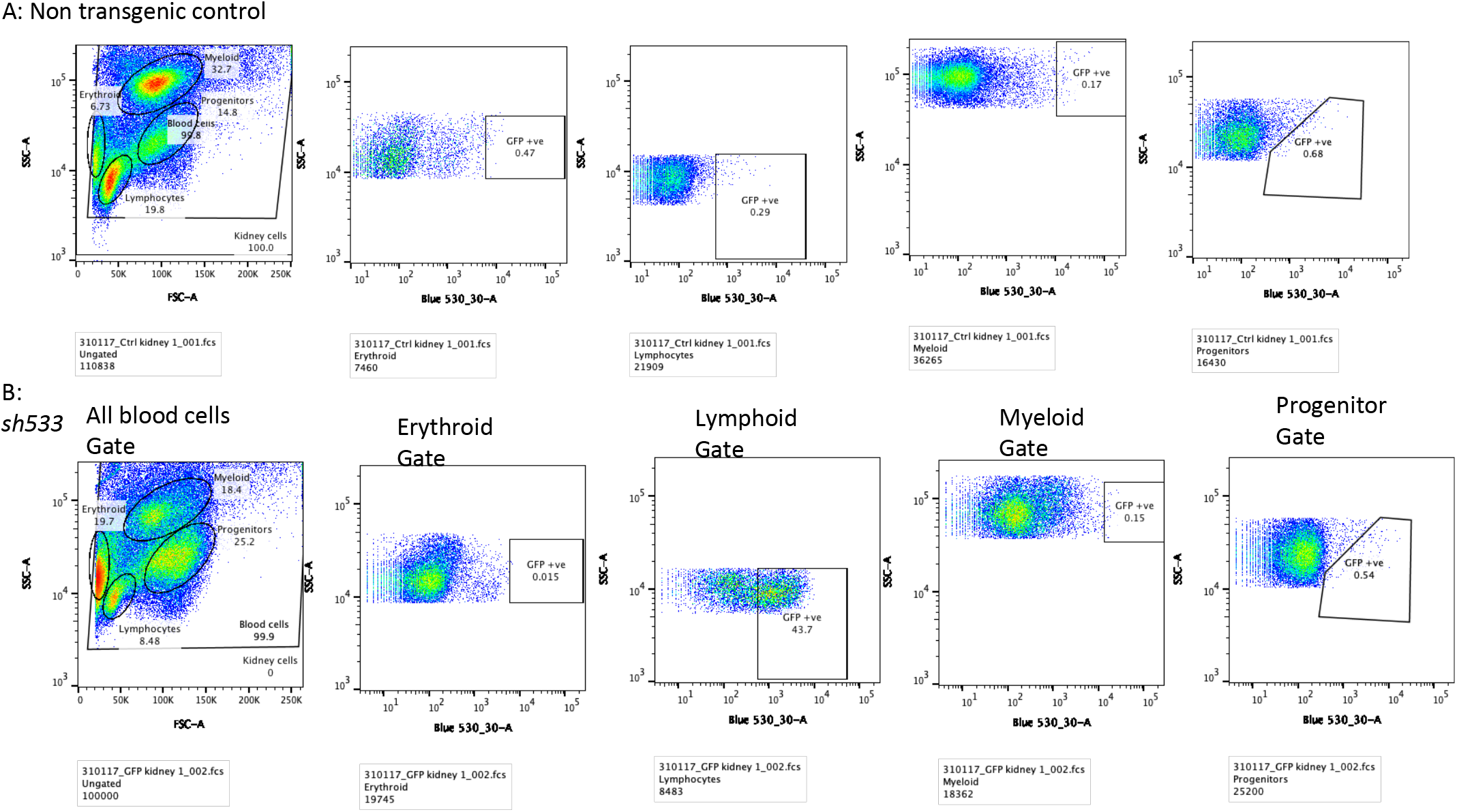
Gating strategy for each heamatopoietic lineages (erythroid, lymphoid, myeloid and progenitor) drawn first in non transgenic control (A) and applied to the sh533 transgenic kidney samples.

**Supplementary Figure 6:**
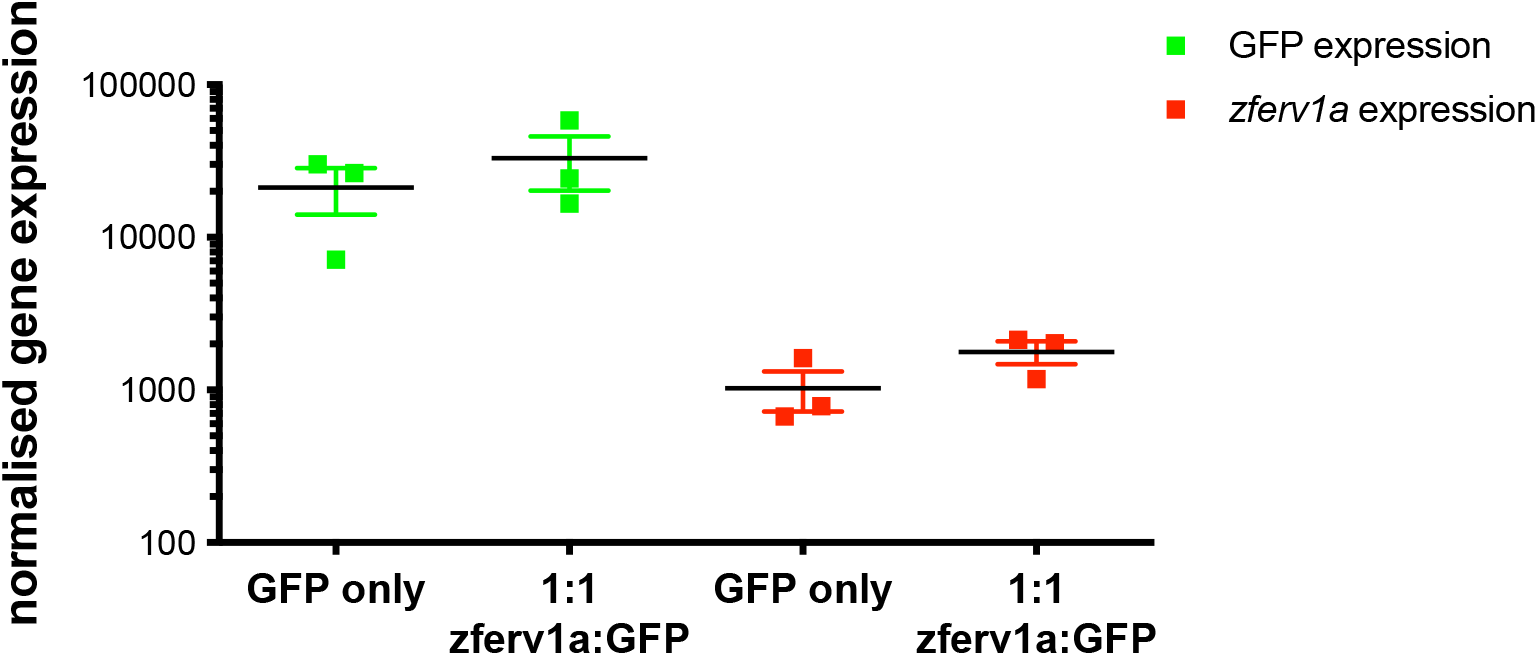
Expression of GFP (green) and zferv1a (red) by qPCR after 1-cell stage injection of a GFP expressing vector alone (GFP only) or in equimolar concentration with zferv1a expression vector (1:1 zferv1a:GFP).

## Notes

### Competing Interest Statement

The authors have declared no competing interest.

### Summary of Updates

We have provided a better characterisation of our novel transgenic line following reviewers comments, alongside a revised analysis of single cell RNA seq and a more complete expression pattern by in situ analysis. The main message of the paper has not changed.

